# Targeting collagen biosynthesis by small molecules inhibiting the function of the peptide-substrate-binding domain of collagen prolyl 4-hydroxylases

**DOI:** 10.64898/2026.02.05.703963

**Authors:** Bukunmi Adediran, Carlos Vela-Rodríguez, Sudarshan Murthy, M. Mubinur Rahman, Rik K. Wierenga, Lari Lehtiö, M. Kristian Koski

**Affiliations:** Faculty of Biochemistry and Molecular Medicine and Biocenter Oulu, University of Oulu, Oulu, Finland

**Author notes:** All authors contributed equally (Bukunmi Adediran, Carlos Vela-Rodríguez, Sudarshan Murthy).

**Keywords:** Peptidomimetics, fibrotic diseases, FRET, High-throughput screening, drug discovery

## Abstract

Collagen prolyl 4-hydroxylase (C-P4H) is an essential enzyme in collagen synthesis and known to be a potential target for drugs that prevent excess collagen formation. Currently known C-P4H inhibitors target the catalytic site of C-P4H, being analogues of 2-oxoglutarate (2OG). However, in mammalian cells there are many other 2OG-dependent dioxygenases with a highly similar catalytic domain, which limits the selective specificity of drugs targeting C-P4H activity. The peptide-substrate-binding (PSB) domain is unique for the C-P4H family and known to be important for the catalytic efficiency of C-P4H. Therefore, interfering with peptide binding to the PSB domain might allow more specific inhibition of the hydroxylation activity of C-P4Hs. We developed a robust FRET-based high-throughput screening assay (Z’ > 0.73) based on PSB-peptide interactions. This assay was used to screen a peptidomimetic library of 15614 compounds with the PSB domains of C-P4H isoforms I and II. A hit compound (OUL-PSBi-001) was identified with IC_50_s of 40 µM and 62 µM for PSB-I and PSB-II, respectively. We also showed that this compound indeed inhibits the catalytic activity of the full-length CP4H-I and -II. This FRET assay provides a new strategy for finding selective inhibitors for the treatment of fibrotic diseases and cancer.

## Introduction

Collagens are extracellular matrix (ECM) proteins, which provide mechanical support, participate in cellular development and communication, and are important for tissue repair and wound healing [1]. Common for all collagen polypeptide chains is a region that has repeating X-Y-Gly sequence triplets, which allows the oligomerization of collagens into the typical triple helical structure [1,2]. The proline 4-hydroxylation at position Y is catalyzed by enzymes called collagen prolyl 4-hydroxylases (C-P4Hs) [3–5]. Three C-P4H isoforms in humans have been identified, which are referred to as C-P4H-I, -II, and -III [3]. C-P4Hs require 2-oxoglutarate (2OG), iron (Fe^2+^), molecular oxygen (O_2_) for its prolyl hydroxylase activity, whereas ascorbic acid (vitamin C) is required for being able to catalyze multiple hydroxylation reactions of procollagen [6,7]. C-P4H-I is the most abundant and most important isoform, whereas C-P4H-II is more tissue and cell type specific [8,9]. During embryonic development collagen synthesis has a critical role, and C-P4H-I knock-out mice have been shown to be embryonically lethal [10]. In healthy adults, collagen synthesis is less critical, although prolonged C-P4H deficiency, due to the lack of vitamin C, leads to scurvy disease [5]. Excessive C-P4H activity is associated with fibrosis caused by the accumulation of collagens [11]. Fibrosis is a major health issue, and an effective way to prevent collagen accumulation is to inhibit collagen synthesis by inactivating the C-P4Hs [5,6,12]. Cancer metastasis is correlated with significant changes of the ECM properties, associated with increased collagen synthesis [13,14], and it has been reported that inhibition of C-P4H-I and C-P4H-II activity dramatically decreases spontaneous metastasis in many cancers [14–16].

C-P4Hs in vertebrates are α_2_β_2_ heterotetramers of about 240 kDa, consisting of two catalytic α- subunits and two protein disulfide isomerases (PDI), which act as β-subunits (**Figure 1a**) [3,5]. The C-terminal half of the α-subunit contains the key residues for Fe^2+^and 2OG binding and is referred to as the catalytic (CAT) domain, as it is responsible for catalyzing the hydroxylation process [17,18]. The CAT domain interacts tightly with three thioredoxin domains of the β- subunit, namely with domains **a, b’** and **a’** [18] (**Figure 1a**). The N-terminal half of the α-subunit contains two helical domains referred to as the N-terminal dimerisation domain and the middle peptide-substrate-binding domain (PSB). In the current structural model of the α_2_β_2_ tetramer, the PSB domain does not interact with the β-subunit but instead protrudes out of the elongated tetrameric molecule, which is capped by the β-subunits (**Figure 1a**) [18].

**Figure 1.**
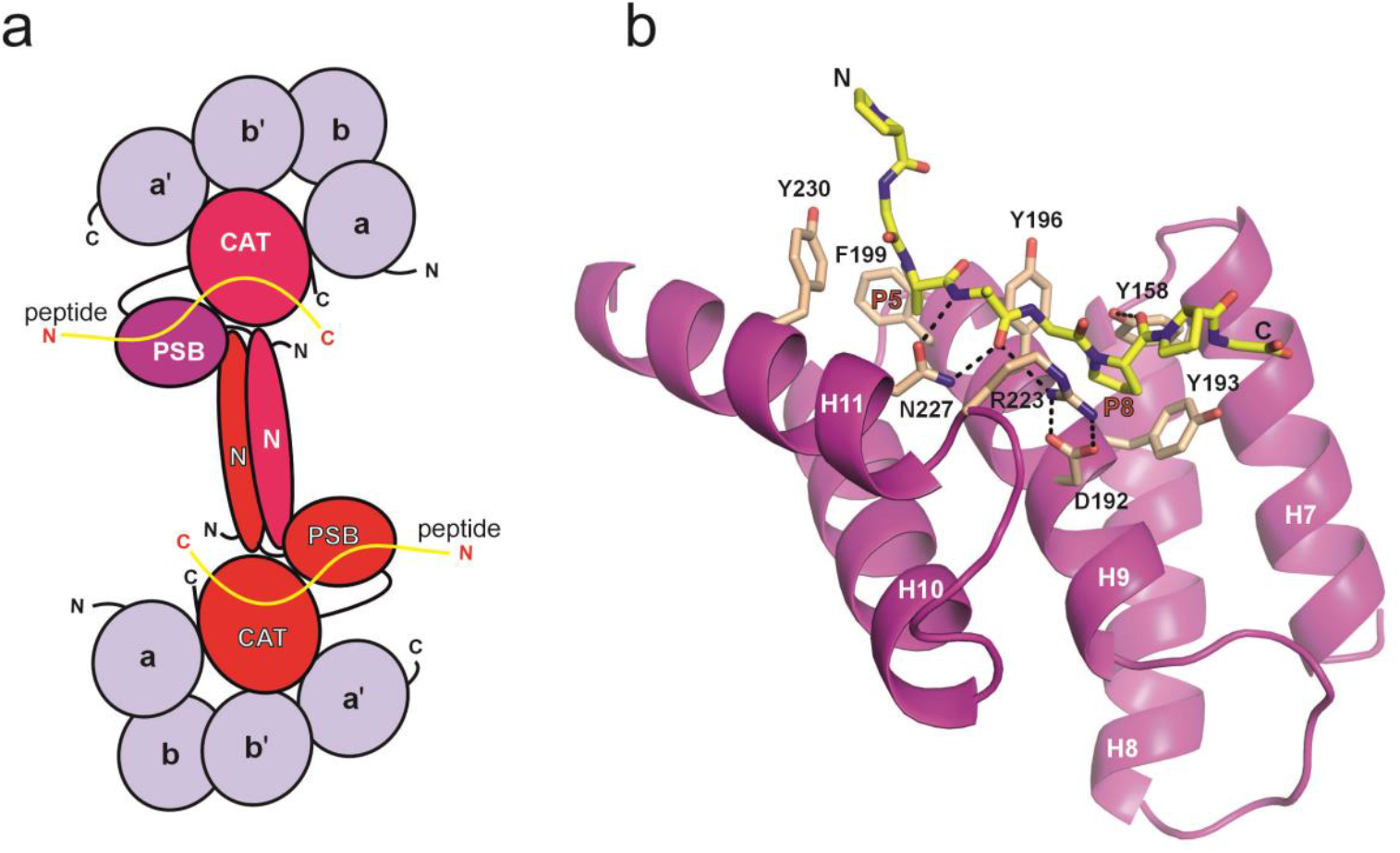
Organization of the C-P4H complex. (**a**) The C-P4H tetramer is an elongated molecule, where the α_2_-dimer (monomers in magenta and red) forms the core of the complex via its dimerisation domains (or N domain), and the CAT domains of the α-subunit interact with the PDI/β-subunits (light blue), which consist of four domains **a, b, b’** and **a’** [18]. The N, PSB, and CAT domains of the “upper” chain of the α_2_-dimer is colored in magenta, purple and magenta, respectively, whereas the domains of the “lower” α chain are red. The PSB domains are protruding out of the core of the tetramer complex but interact with the dimerisation (N) domain and the CAT domain. The proposed binding mode of the procollagen peptide (yellow) to the PSB and CAT domains is also indicated. (**b**) Crystal structure of PSB-II with bound PAG peptide (PDB ID 6EVN) [25]. The PSB domain is composed of five α-helices H7-H11. The PAG peptide binds to this domain in such a way that the proline of the central Pro-Ala-Gly triplet occupies the P5 pocket (lined by Tyr196, Phe199 and Tyr230), whereas the first proline of the following Pro-Pro-Gly triplet is buried in the P8 pocket (lined by Tyr158, Tyr193, Tyr196, and the salt bridge Asp192-Arg223). The protein-peptide interactions concern mainly the stacking interactions between tyrosine side chains of the domain and prolines of the peptide. The side chains of Asn227, Arg223 and Tyr158 form the key protein-peptide hydrogen bonds (as visualized by dotted lines).

The three C-P4H isoforms all have different α-subunits, each encoded by its own gene [19,20]. The α-subunit of each isoform contains the PSB domain which consists of five α-helices (**Figure 1b**) and it is thought to be important in anchoring the long procollagen chain during its hydroxylation [18,21–24]. In the current C-P4H model, the peptide binding sites of the PSB and CAT domains are properly aligned for simultaneous binding to an extended peptide (**Figure 1a**) [18]. It has been shown that tyrosine-to-alanine point-mutations of the tyrosines that line the peptide-substrate binding groove of the PSB domain (especially tyrosines 193, 196 and 230, see **Figure 1b**) lower the activity of the full-length C-P4H-I for the proline-rich model peptide substrate (PPG)_10_ indicating that the PSB domain is important for efficient proline hydroxylation by the CAT domain [23]. The PSB domains of C-P4H-I and C-P4H-II (henceforth referred to as PSB-I and PSB-II, respectively), have a sequence identity of 65%, and they have somewhat different peptide binding specificity and properties [21,24–26]. Crystal structures of both PSB-I and PSB-II in apo form and in complex with short proline-rich peptide substrates have been reported (**Figure 1b**) [23–26]. Both PSB domains have been co-crystallized with a procollagen-like peptide PPG-PAG-PPG (also referred to as the PAG peptide), which has the binding affinity (*K*_D_) to PSB-I and PSB-II of 91 µM and 20 µM, respectively [25,26].

Since its discovery, C-P4H has been thought to be a possible pharmaceutical target due to its role in promoting fibrosis. The identified inhibitors so far described in the literature are metal ions which replace the active site Fe^2+^, or cofactor analogs which replace 2OG in the active site of the CAT domain [3,5,6]. However, because C-P4Hs share a common catalytic domain with other P4H homologs and other 2OG dependent dioxygenases [27,28], designing specific and selective inhibitors for the CAT domain of C-P4Hs is challenging. Currently there are no C-P4H inhibiting drugs in the market.

We rationalized that a feasible alternative approach to inhibit C-P4H activity could be to inhibit the binding of procollagen to the PSB domain as this domain is specific for C-P4Hs and important for its catalytic function. Herein, we present high-throughput (HTS) screening assay for finding inhibitors of the PSB domain function based on förster resonance energy transfer (FRET), which has been used successfully to probe protein-protein interactions of other complexes [29]. After screening a library of 15614 peptidomimetic compounds with PSB-I and PSB-II, we identified one potential compound which binds to the PSB domain of both C-P4H orthologues. Our findings support the concept that inhibitors of PSB peptide binding are also inhibitors of the C-P4H catalytic activity. The discovered compound is the first reported C-P4H inhibitor targeting the C-P4H PSB domain.

## Results

### CFP-tagged PSB-I and PSB-II bind to the YFP-PAG peptide resulting in FRET signals

As a starting point for developing the FRET screening assay targeted to find binders for the PSB domain, it was known from the previous studies that PSB-I and PSB-II domains bind differently to polyproline peptides and peptides containing only PPG triplets [21–25]. However, it was shown that both constructs bind the PAG peptide by using the same mode of binding and with affinity better than 100 µM [25,26], which would allow making a screening assay suitable for both isoforms. The screening assay development was started with the PSB-II domain. The N-termini of the PSB-II domain and the PAG peptide were fused to the C-terminus of the monomeric variants of CFP and YFP, respectively (**Figure 2**). This fusion allows for the use of FRET to monitor and analyze peptide binding. When the CFP-fused protein and the YFP-fused peptide form a complex, the proximity of the fluorophores facilitates energy transfer from CFP to YFP. The presence of an inhibitor of peptide binding disrupts this complex formation, leading to a reduction in energy transfer (**Figure 2**). This inhibition of peptide binding is manifested therefore as a higher emission at the CFP emission maximum wavelength (477 nm) and a lower emission at the YFP emission maximum wavelength (527 nm). To quantify the amount of energy transferred from CFP-PSB-II to YFP-PAG when present in the complex, the ratiometric FRET value (rFRET) is used. This rFRET value is obtained by dividing the fluorescence measured at 527 nm by the fluorescence measured at 477 nm. A lower rFRET value indicates that a lower amount of the complex is present in the reaction mixture, because of the presence of an inhibiting small molecule (**Figure 2**).

**Figure 2.**
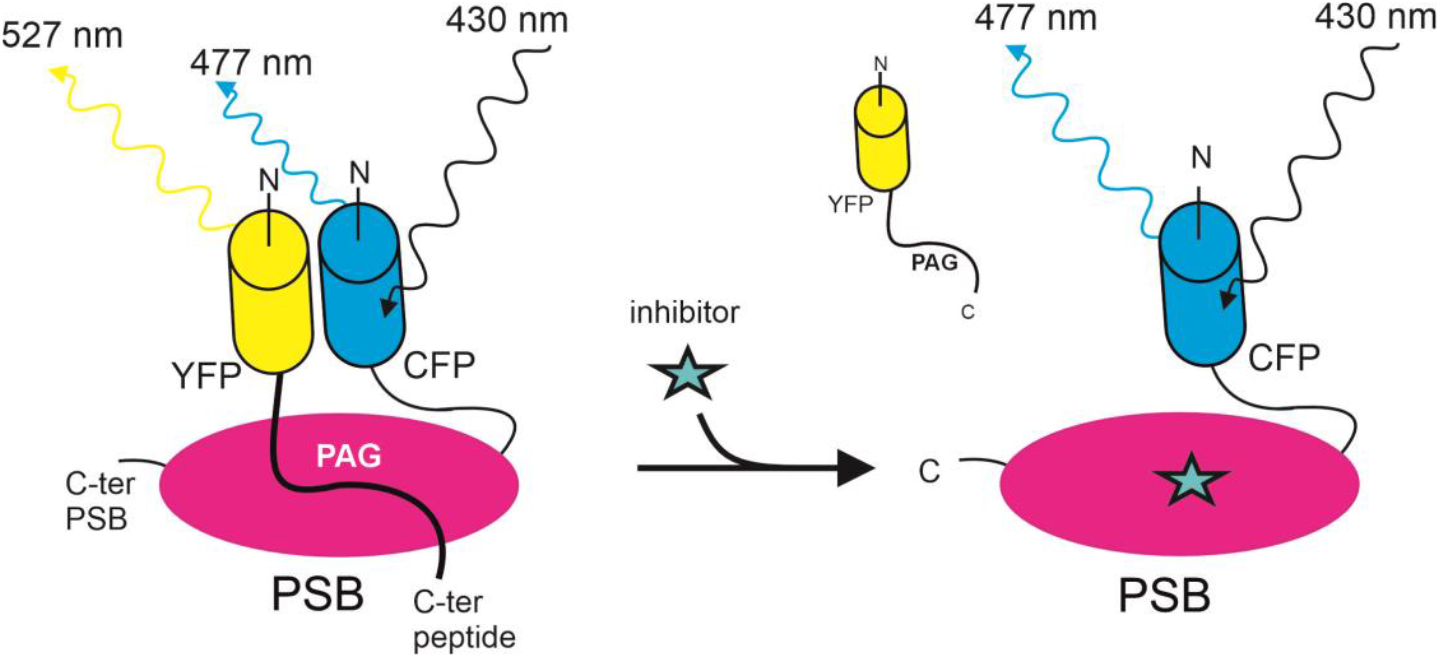
Schematic representation of the FRET principle: the binding of a small molecule (indicated by a star) disrupts the interaction between the PSB domain (tagged with CFP at its N-terminus) and the PAG peptide (tagged with YFP at its N-terminus). The rFRET value, the ratio of the emission at 527 nm divided by the emission at 477 nm, is low in the presence of an inhibitor.

Assay parameters such as concentration, molar ratio, buffer, and reaction volume were tested to achieve a reproducible and robust signal for the CFP-PSB-II – YFP-PAG pair. The optimized conditions of the assay mixture included 10 mM Bis–Tris-Propane (pH 7.0), 3% (w/v) PEG 20K, 0.01% (v/v) Triton-X100 and 0.5 mM TCEP working buffer, 100 nM CFP-PSB-II concentration, 1:20 CFP:YFP ratio, and reaction volumes ranging 10 to 30 µL. These assay conditions were found to be suitable also for the CFP-tagged PSB-I construct. The statistical fluctuations observed in the measurements for both assays were found to be very similar (**Figures 3a, b**). Based on the Z’ number calculated from the plate validation (0.79 for PSB-I and 0.73 for PSB-II), the assay can be considered excellent for inhibitor screening [30]. Given that compounds are typically dissolved in DMSO, its effect on the assay was investigated by using the PSB-II construct. We observed that the assay can tolerate DMSO up to 5%, which exceeds the percentage of DMSO used in the studies **(Figure 3c)**. Initial experiments utilized free YFP to confirm that the recorded rFRET signal was indeed a consequence of the specific interaction between PSB and the peptide of the PSB-peptide complex, rather than non-specific energy transfer. We have earlier reported for another system that in the presence of 1 M guanidine hydrochloride (GndHCl), the protein-protein interactions are effectively disrupted with little to no effect on fluorescence properties of the stable YFP and CFP tags [31]. Therefore, this condition was used as a positive control in the inhibitor screening.

**Figure 3.**
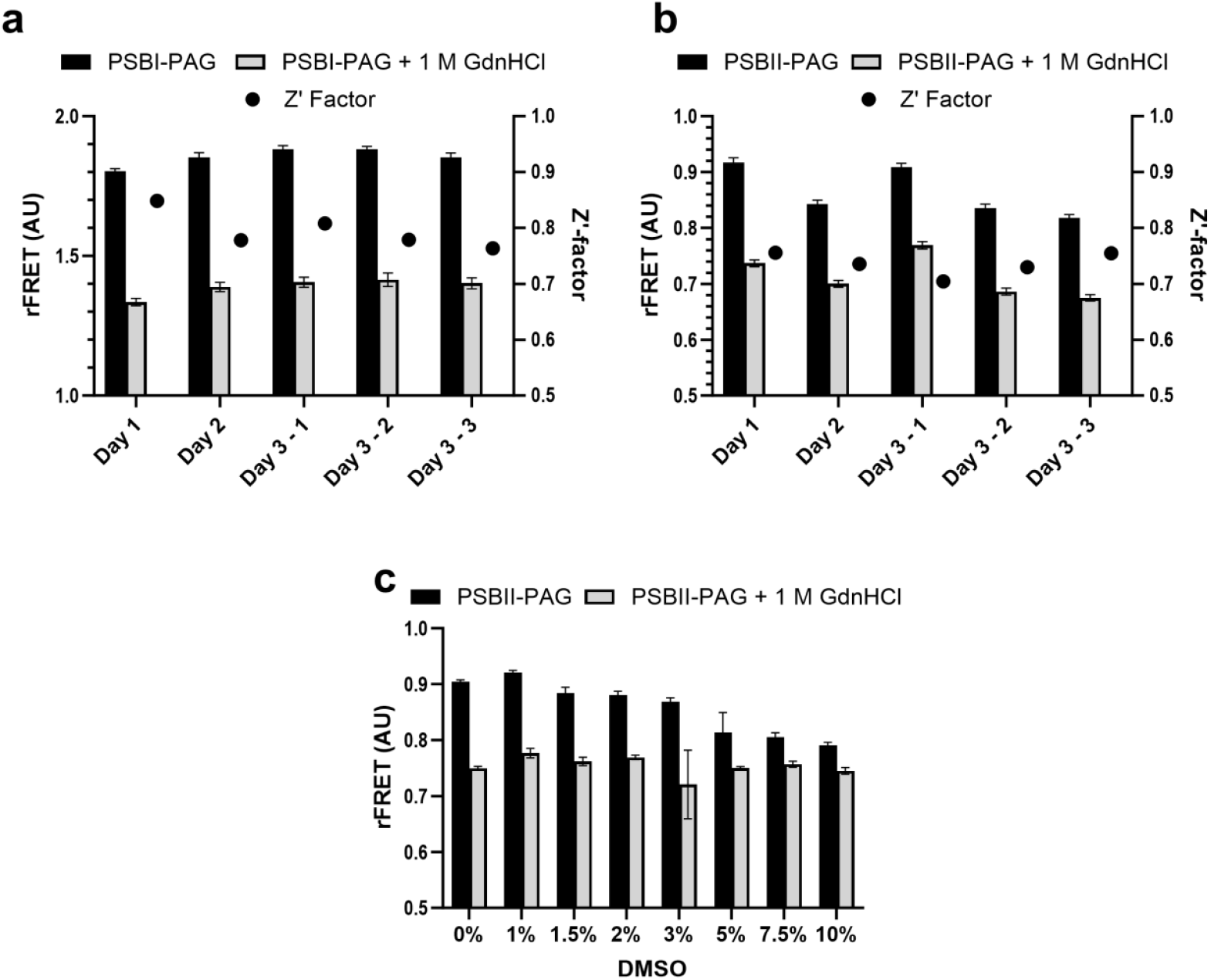
The FRET-based assay is suitable for screening of small molecules. **(a)** Plate validation for CFP-PSB-I. **(b)** Plate validation of CFP-PSB-II. **(c)** DMSO tolerance test of the assay for the CFP-PSB-II / YFP-PAG FRET pair. The reaction buffer supplemented with 1M GdnHCl was used as a positive inhibition control. The data presented in **(a)** and **(b)** corresponds to the mean and SD of 176 replicates per plate. The data in **(c)** corresponds to the mean and SD of four replicates. The concentration of CFP-PSB used was 100 nM and for YFP-PAG was 2 µM.

### Dissociation constants for the PSB:PAG interactions measured with FRET

To validate the assay and confirm that the fluorescent tags did not interfere with peptide binding, the FRET method was used to measure binding affinity of the CFP-tagged PSBs and YFP-tagged PAG peptide. While FRET has been used to determine dissociation constants (*K*_D_) for synthetic fluorophores [32], it has also been adapted for fluorescent proteins [31]. The rFRET value was calculated from the fluorescence data and the saturation curve was plotted with a *K*_D_ values of 12.1 µM and 17.5 μM for PSB-I:PAG and PSB-II:PAG complexes, respectively (**Figure 4**). These confirm that the CFP and YFP tags do not reduce binding affinity and that both PSBs would have similar affinity to the peptide. It can be noted that the previously reported values of PSB-I:PAG and PSB-II:PAG are 92 µM and 19 µM, respectively [25,26].

**Figure 4.**
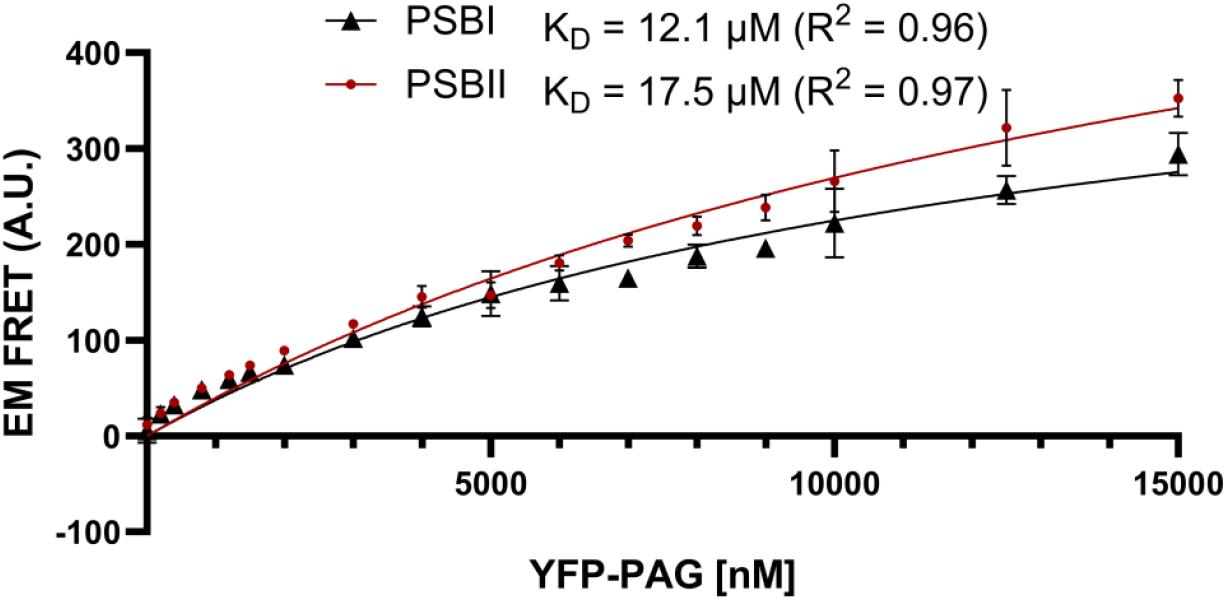
FRET-based saturation curve for binding of the PAG peptide to PSB-I and PSB-II. The measurements show the mean and standard deviation from three independent experiments with 4 technical replicates wherein the concentration of CFP-PSB-I (triangle) and CFP-PSB-II (dot) was kept constant at 200 nM.

### The FRET-based inhibitor screening identifies a compound which binds to both PSB domains with high affinity

A small-molecule library consisting of 15614 peptidomimetic compounds in assay plates provided by the Institute for Molecular Medicine Finland (FIMM) and used for the initial screening step of both PSB-I and PSB-II (**Figures 5a, b**). The used compound concentration in the screen was 20 µM and the DMSO concentration was less than 0.2%. The screening data were analyzed with multiple filters to eliminate non-specific FRET signals. Assay mixtures for which the fluorescence with compounds was more than 20% greater than the fluorescence of the negative control at 527 nm were considered outliers and eliminated. Furthermore, fluorescence of samples with compounds more than 20% greater than the positive control at 477 nm were considered as outliers as well. Also, compounds showing signs of quenched fluorescence were eliminated. After applying these filters, we set the hit limit of the rFRET value to 40% (**Figures 5a, b**). The initial screening yielded 57 positive hits, and these compounds were rescreened subsequently with PSB-I and PSB-II (**Figure 5c**). The 57 hit compounds mostly inhibited both PSBs but there is some clear variation between the inhibition efficacies (**Figure 5c**)

**Figure 5.**
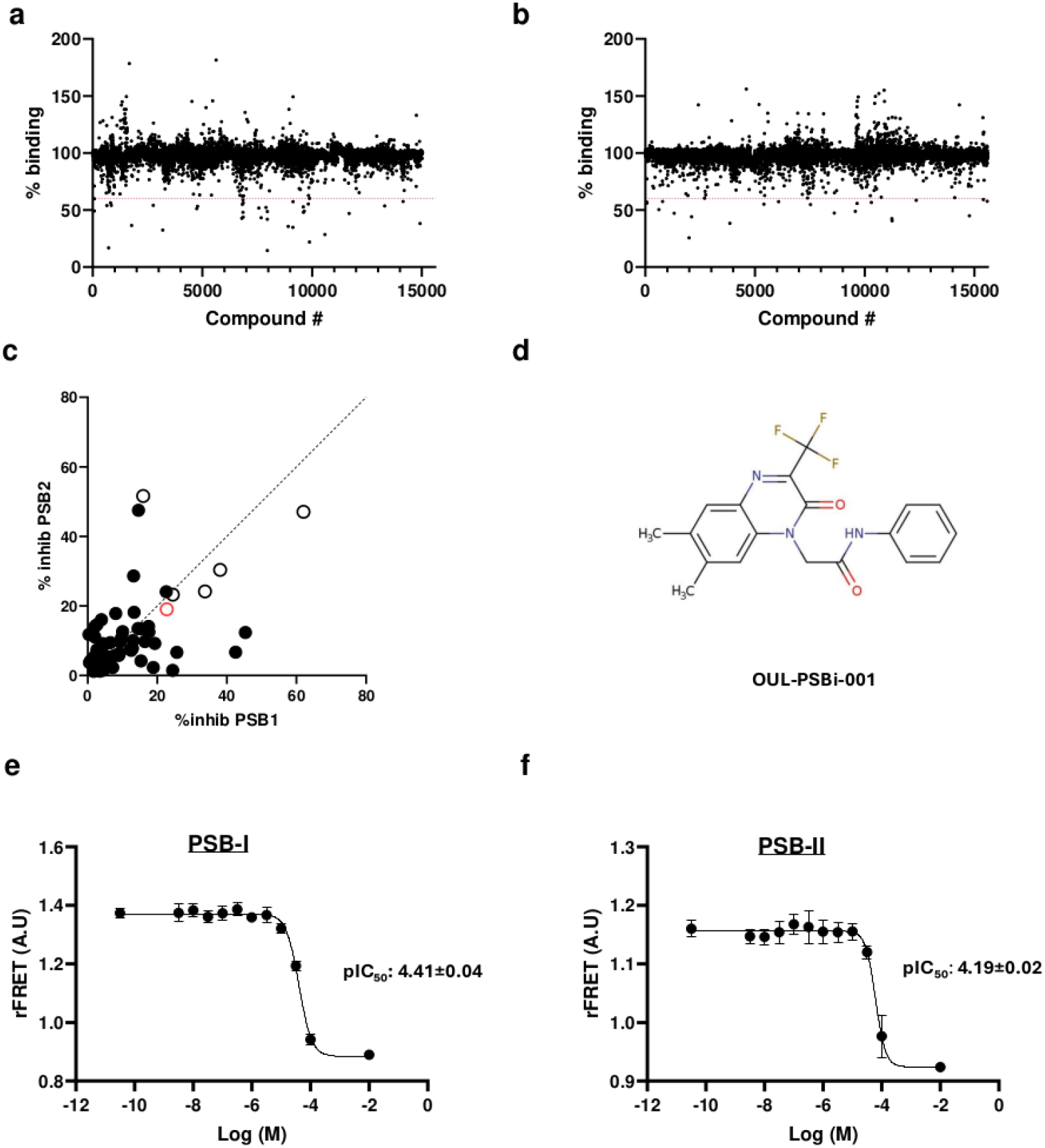
Identification of chemical compounds that inhibit the interaction between PSB and a proline-rich peptide using a FRET-based assay. Screening of the peptidomimetic library for: **(a)** PSB-I and **(b)** PSB-II. The red dotted line represents the inhibition threshold used to select compounds from the primary screening. **(c)** Correlation of the apparent inhibition of compounds selected from **(a)** and **(b)**, plotted with PSB-I on the x-axis and PSB-II on the y-axis. Open circles highlight the compounds which were ordered and studied further. The red open circle corresponds to the OUL-PSBi-001 **(d)** Structure of the hit compound OUL-PSBi-001. The quinoxaline core is on the left side, and the N-phenyl moiety is on the right side. **(e)** Determination of the IC_50_ values of the hit compound against PSB-I, and **(f)** against PSB-II.

To identify hits with significant potency, we purchased the six most potent hit compounds, which were available from commercial vendors, for potency measurements (**Figure 5c**). The best hit was compound called 2-[6,7-dimethyl-2-oxo-3-(trifluoromethyl)-1,2-dihydroquinoxalin-1-yl]-N-phenylacetamide (**Figure 5d**, referred to as OUL-PSBi-001 from now on), which showed full inhibition at 100 µM. The measured IC_50_ for PSB-I was 40 µM and for PSB-II 62 µM (**Figure 5e**). The binding of OUL-PSBi-001 to the PSB domains was also validated with the nano differential scanning fluorimetry (NanoDSF) by using the PSB domains without the CFP tag. Both PSB isoforms showed higher melting temperatures in the presence of 100 µM OUL-PSBi-001 in the solution. The T_m_ of the PSB-I domain increased from 63.8°C to 64.2°C, and the T_m_ of the PSB-II domain increased from 62.4°C to 62.9°C. The increased thermal stabilities are consistent with binding of this compound to the PSB domains.

### OUL-PSBi-001 inhibits the catalytic hydroxylase activity of C-P4H-I and C-P4H-II

To measure the effect of OUL-PSBi-001 on the hydroxylase activity of C-P4H-I and C-P4H-II, we established a microplate-based bioluminescence assay using full-length human C-P4H-I and C-P4H-II, purified from an insect-cell expression system [33]. The assay is the same as used previously for measuring the catalytic activity of lysyl hydroxylase enzymes [34]. In this assay, succinate formation is monitored with the Succinate-Glo™ JmjC Demethylase/Hydroxylase Assay kit (Promega, Madison, WI; Cat# V7990), which converts succinate generated by the C-P4H hydroxylase reaction into a luminescent signal (**Figure 6a**). The proline-rich peptide (PPG)_10_ was used as the substrate, as it is a well-characterized C-P4H substrate [9]. The assay mixture including C-P4H-I (5 µg/mL) or C-P4H-II (5 µg/mL) and peptide substrate (100 µM) produced a high luminescent signal after 30 min reaction time (**Figures 6b, c**). Including 100 µM OUL-PSBi-001 in the reaction mixture reduces the background corrected signal for both C-P4H-I as well as C-P4H-II by about 50% **(Figures 6b, c)** showing that OUL-PSBi-001 inhibits the activity of both full-length C-P4H isoforms.

**Figure 6.**
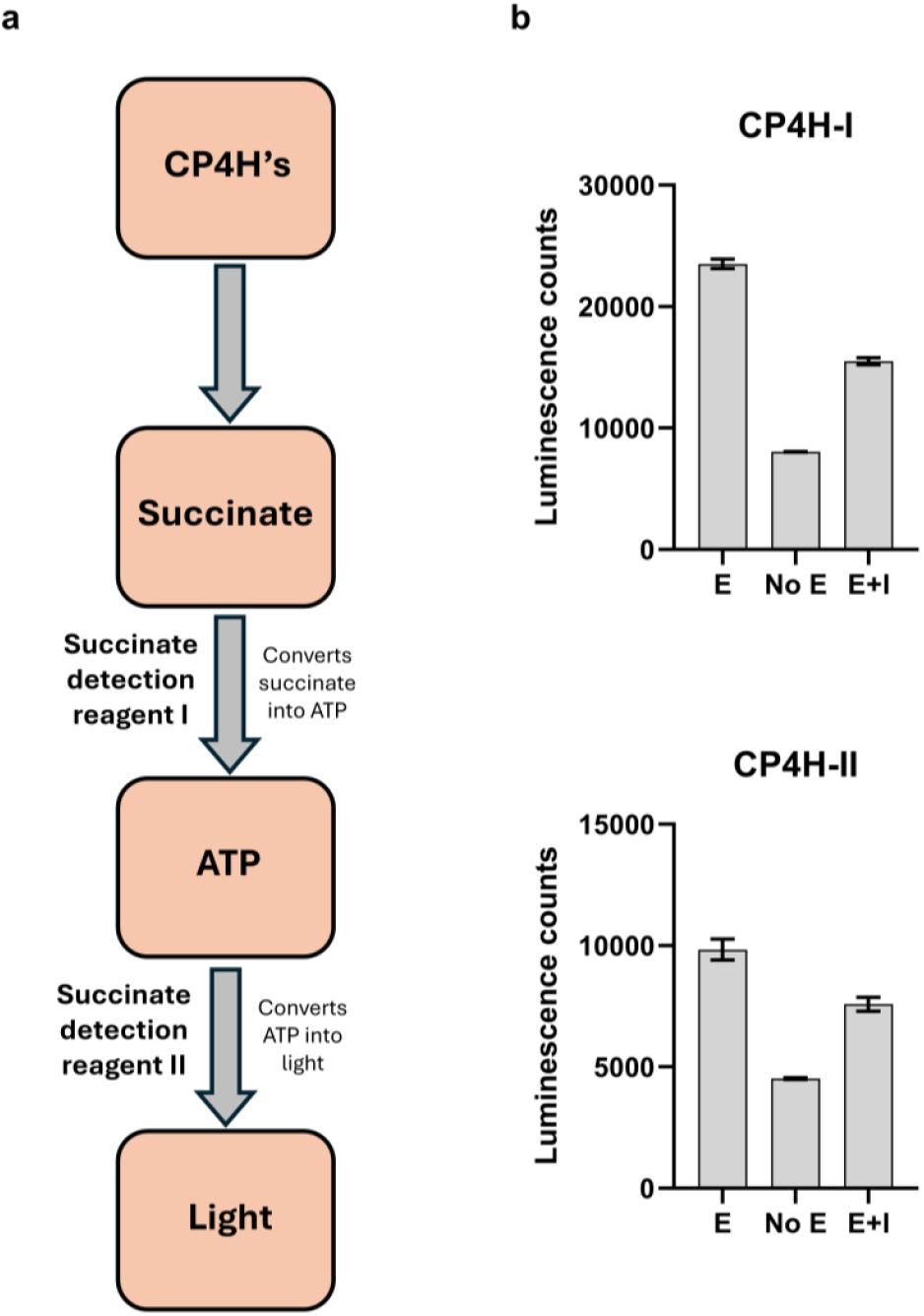
Inhibition of full-length C-P4Hs with OUL-PSBi-001 measured using a bioluminescence-based C-P4H activity assay. **(a)** Schematic overview of the assay principle. **(b)** Activity assay results of C-P4H-I (upper panel) and C-P4H-II (lower panel) showing the uninhibited enzyme reaction (“E”), the control reaction not including any enzyme (“No E”), and the inhibited reaction including enzyme and 100 µM OUL-PSBi-001 (“E+I”).

### Docking studies show that the OUL-PSBi-001 compound fits in the peptide-binding groove of the PSB domain

Docking studies were performed using the crystal structures of PSB-I (PDB ID 9HT8) and PSB-II (PDB ID 6EVN) complexed with the PAG peptide [25,26]. The docking results propose that the compound binds in the peptide-binding groove of both PSB isoforms in such a way that the quinoxaline core binds to the P5 pocket and the N-phenyl moiety occupies the P8 pocket (**Figures 7a, c**). The two methyl substituents of the quinoxaline moiety point inwards and have van der Waals interactions with the protein and the trifluoro methyl group points to the solvent region. Superimpositions of the OUL-PSBi-001 docking models with the crystal structures of the PAG-complexed PSB-I (**Figure 7b**) and PSB-II (**Figure 7d**) show that the quinoxaline core aligns well with the peptidyl proline bound to the P5 pocket, whereas the N-phenyl group aligns well with the peptidyl proline bound in the P8 pocket. In the docking models of both PSB isoforms, two direct hydrogen bonds are found between, respectively, the side chains of Asn227 and Arg223 and the compound carbonyl oxygen (**Figures 7b, d**). These hydrogen bonds correspond to the direct hydrogen bonds seen in the crystal structures of the PSB-peptide complexes. In these complexes, the side chains of Asn227 and Arg223 are hydrogen bonded to the back-bone carbonyl oxygen of the alanine of the PAG peptide. The other critically important hydrogen bond found in the PSB-peptide complexes, between the Asn227 side chain and the backbone peptide NH of the alanine of the PAG peptide **(Figure 7)**, is not found in the docking models of PSB and OUL-PSBi-001 due to the absence of a hydrogen bond donor (corresponding to the free peptide NH of the alanine residue of the bound peptide) in the docked compound.

**Figure 7.**
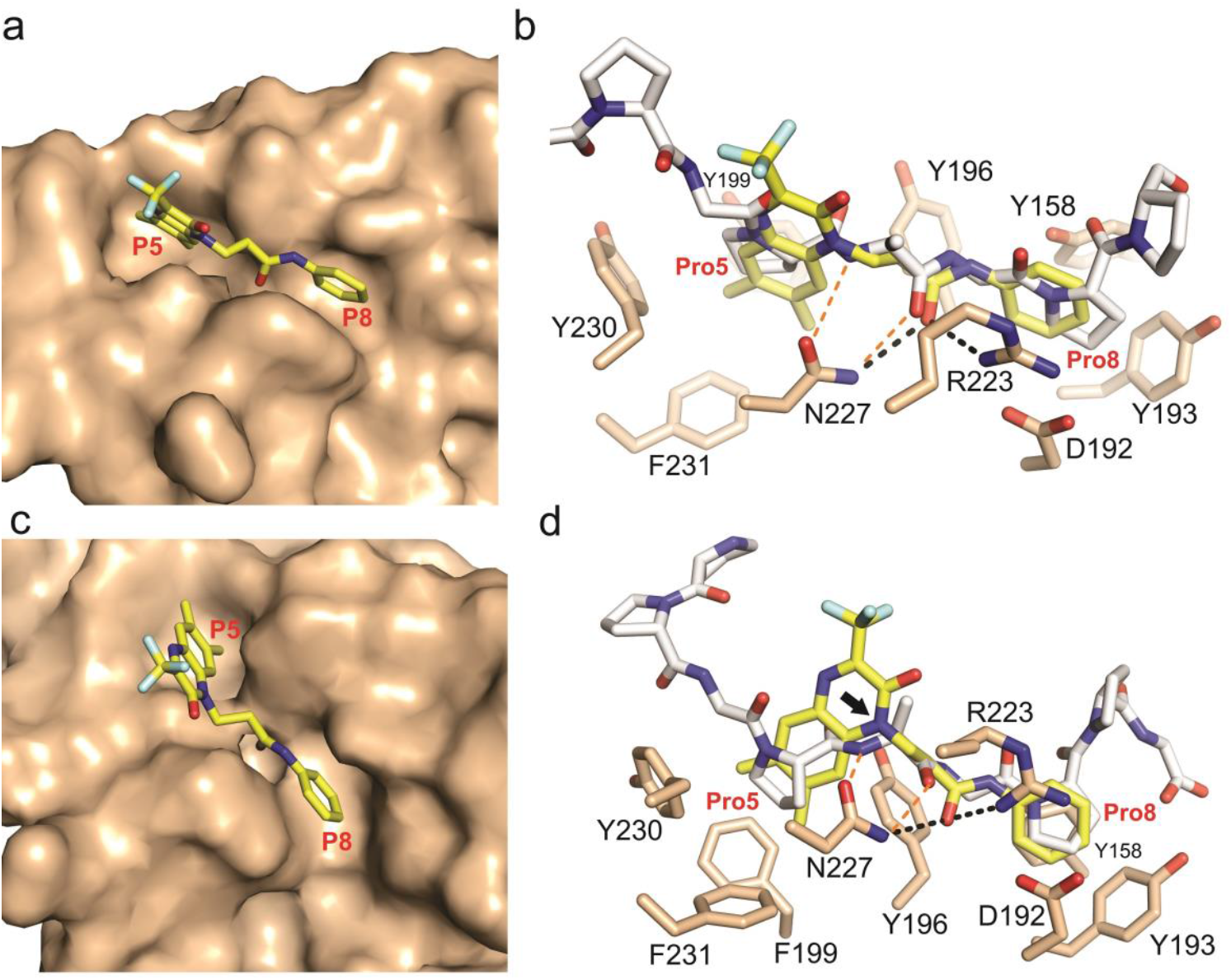
Docking models of compound OUL-PSBi-001 with the PSB domains. **(a)** The OUL-PSBi-001 compound (yellow sticks) modelled in the peptide binding groove of PSB-I (light brown surface) in such a way that the quinoxaline moiety (left part) and the N-phenyl group (right part) of this compound occupy the P5 and P8 pockets, respectively. **(b)** The proposed mode of binding of the OUL-PSBi-001 compound mimics the binding of the PAG-peptide (white sticks, taken from the PSB-I PAG complex structure, (PDB ID 9HT8). The model also proposes two direct hydrogen bonds between the carbonyl oxygen of the inhibitor and the PSB domain as highlighted in the figure with the black dashed lines. **(c)** The docking model of OUL-PSBi-001 with PSB-II is highly similar to the PSB-I docking model. **(d)** The proposed hydrogen bond interactions between PSB-II and OUL-PSBi-001 (black dotted lines) are the same as in the corresponding complex of PSB-I and OUL-PSBi-001 (panel **b**). This view (slightly rotated compared to panel **b**) shows that the second amide moiety of the compound (black arrow) is not free to form a hydrogen bond with Asn227 like seen in the peptide complexes. Also highlighted in panels **(b)** and **(d)** are the hydrogen bond interactions (by orange dotted lines) between the alanine of the peptide and the side chain of Asn227 of PSB.

## Discussion

C-P4Hs are long-known potential targets for treatment of fibrotic diseases [12] and more recently also for treatment of cancer progression [14]. Many 2OG analogues (e.g. pyridine 2,4-dicarboxylate, 3-hydroxypyridine-2-carbonyl-glycine, oxalylglycine, 3-carboxy-4-oxo-3,4-dihydro-1,10-phenanthroline) and metal ions (e.g. Zn^2+^, Co^2+^) are known C-P4H inhibitors. These compounds bind, respectively, to the 2OG and iron binding sites of the CAT domain, and prevent the hydroxylation reaction, but not the peptide binding [6]. These compounds also inhibit other mammalian P4Hs like hypoxia inducible factor (HIF) prolyl 4-hydroxylase (HIF-P4H, also referred to as PHD2) [6], which have a highly homologous CAT domain [35]. Inhibitors against HIF-P4H have been studied intensively during the past twenty years. The first inhibitor, a 2OG analogue Roxadustat/Evrenzo, has been accepted for treatment of renal anemia in several countries [36,37]. Roxadustat has also been shown to inhibit C-P4Hs with similar efficacy [38]. In addition, there are several other 2OG-dependent dioxygenases in human cells with a similar CAT domain fold [27], and inhibitors targeted to the C-P4H CAT domain can also reduce the activity of these other enzymes. There is therefore an urgent need to develop new strategies for finding inhibitors of the C-P4H enzymes and targeting their PSB domain would be an excellent approach. The PSB domain is not found in HIF-P4H, and other 2OG-dependent dioxygenases and its peptide binding properties have been shown to be important for the catalytic activity of C-P4H. In particular it was found that point mutation variants of C-P4H-I in which tyrosine residues of the P5 and P8 pocket were mutated into an alanine have *K*_M_ values of the model substrate (PPG)_10_ that are increased from 30 µM for wild type C-P4H-I to about 250 µM [23]. The precise role of the PSB domain in the reaction mechanism of C-P4H is not understood, but it is thought to act by providing an anchoring binding site for the procollagen chain during hydroxylation by the catalytic domain [18]. This information has provided the basis for our new approach aimed at finding, ultimately, drugs that inhibit the C-P4H enzymes by binding to the PSB domain.

For this approach, we developed a new FRET-based HTS platform to identify compounds that bind to the PSB domain. The isolated PSB domains are suitable for HTS as they can be produced efficiently recombinantly in the *E. coli* host, and they are small and stable proteins [21–23]. Furthermore, our extensive structural and biophysical peptide-binding studies have shown that the PAG peptide having the PxGP motif binds to both PSB domains in a similar way and with similar affinity [25,26]. The YFP-tagged PAG peptide can be produced in a straightforward manner in *E. coli* and therefore this tagged peptide is an optimal component in this screening method with the CFP-tagged PSB-I and PSB-II isoforms. The assay was optimized and validated leading to the generation of a highly robust and reproducible signal with a Z’-factor >0.73.

In contrast to other developed inhibitor screening assays of C-P4H-I based on the quantification of succinate produced by the hydroxylation reaction [39], this FRET method directly quantifies the interaction of the PSB domain and the procollagen peptide. Therefore, the identified hit is expected to inhibit the protein-peptide interaction and not the catalytic activity of the CAT domain of C-P4H. Indeed, our results confirm that the hit compound OUL-PSBi-001 inhibits the C-P4H reaction (**Figure 6b**). The hit compound, OUL-PSBi-001 (PubChem CID: 50760839) is a peptidomimetic with two amide bonds and a quinoxaline core (**Figure 5d**). Quinoxaline-based analogues are known drug molecules. They are used to inhibit tyrosine kinases in cancer therapy [40] and known to bind to the ATP-binding pocket of the kinases as shown by the crystal structure of JAK2 [41].

Our studies show that the developed FRET assay can identify small molecule inhibitors of the formation of PSB-peptide complexes from a large library of compounds and that it can be used with the two most abundant PSB isoforms (PSB-I and PSB-II) for the discovery of compounds that inhibit the PSB function. The identified hit compound provides the first step in the development of potent PSB binders. Structural studies of the PSB domains complexed with this inhibitor will provide more insight into its binding mode and allow for a structure-based design of tighter binding analogs. As the compound also shows inhibition of the C-P4H enzymatic activity with the peptide (PPG)_10_ as substrate, it also allows for mechanistic studies of how the two C-P4H activities, peptide binding by the PSB domain and hydroxylation activity by the CAT domain, are acting in concert to hydroxylate procollagen.

## METHODS

### Cloning

Plasmids containing the α-subunits of human C-P4H-I and -II [18], were used as template for the generation of amplicons coding for the PSB domains. Amplicons were then cloned by sequence and ligation independent cloning (SLIC) to a linearized pNIC-CFP vector [31]. The generation of the YFP-PAG peptide was done by amplifying the mCitrine sequence with a 3’ extension coding for the desired amino acid sequence; the amplicon was subsequently cloned by SLIC to a linearized pNIC28-BSA4 vector. Both vectors contain N-terminal 6x His tags for purification purposes. The SLIC reaction was done with a 1:3 ratio transforming the product to XL-1 blue competent cells, which were grown overnight at 37°C on LB agar plates containing 5% sucrose and 50 µg/mL kanamycin. Plasmids were extracted from a single colony, and the cloning was confirmed by sequencing in the Biocenter Oulu sequencing core facility.

### Protein Expression and Purification

The fusion proteins were expressed in *E. coli* BL21(DE3) in 1 L of terrific broth-base auto-induction media supplemented with glycerol (8 % w/v). The cultures were incubated at 37°C and upon reaching an OD600 value of 0.8 the temperature was decreased to 18°C for overnight incubation. Cells were harvested by centrifugation at 5020 ×*g* and re-suspending in 15 mL of lysis buffer [50 mM HEPES (pH 7.5), 500 mM NaCl, 10 % v/v glycerol, 0.5 mM TCEP, 10 mM imidazole]. The re-suspended cells were flash frozen in liquid nitrogen and stored at -20°C until purified.

The fusion proteins were purified using a 5 mL Hi-Trap IMAC HP column (Cytiva) charged with NiSO_4_ and equilibrated with a 20 mL lysis buffer. Proteins were eluted with elution buffer (lysis buffer with 300 mM imidazole). YFP-PAG and CFP-PSB-I eluted as pure protein, as judged by SDS-PAGE, whereas CFP-PSB-II was further purified with size-exclusion chromatography (SEC). The fusion proteins were flash frozen and stored at-70 °C.

### FRET-assay development and optimisation

The FRET assay herein presented was developed using a previously validated approach [31], in which the interaction of two binding partners bring the fluorescent proteins to proximity thus promoting the transfer of energy from CFP to YFP (**Figure 2**).

For the assay, the fluorescence intensities at 477 nm and 527 nm were measured with an Infinite Pro M1000 microplate reader (Tecan) using an excitation 430 nm as the excitation wavelength. The rFRET value was calculated by dividing the intensity at 527 nm by the intensity at 477 nm.

Initially, we determined the effect of the ratio between CFP-PSB-II and YFP-PAG based on previous reports of this interaction [25]. For this, we maintained CFP-PSB-II at 0.5 µM while varying the concentration of YFP-PAG in ratios of 1:2, 1:5, 1:10, 1:15 and 1:20 (PSB:PAG) in a total volume of 30 µL in 384-well flat bottom polypropylen plates (Fisherbrand). YFP lacking the PAG-peptide was used to assess whether rFRET value was specific to the PSB:PAG interaction or otherwise caused by unspecific interaction of the fluorophores at high concentrations. We subsequently optimised the protein concentration by determining the rFRET value as previously while varying the CFP concentration (25, 50, 100, 200, 400 and 500 nM) at a constant 1:20 ratio. As a final parameter before proceeding with the compound screening, we measured the rFRET value with reactions containing 100 nM CFP-PSB-II (1:20 CF:YFP ratio) in 10, 20 and 30 µL. For all stages of the optimisation we used 10 mM Bis–Tris-Propane (pH 7.0), 3% (w/v) PEG20K, 0.01%(v/v) Triton-X100 and 0.5 mM TCEP as the reaction buffer.

### Dissociation constant determination

To determine the dissociation constant of the CFP-PSB-II:YFP-PAG pair, the concentration of the CFP-PSB-II was kept constant (200 nM) while the concentration of YFP-PAG was gradually increased from 0 to 15 uM. 10 uL reaction volume and 4 replicates were used per concentration condition. Each sample was excited at 430 nm and the emission fluorescence was measured at 477 nm and 527 nm. In addition, samples were further excited at 477 nm and emission fluorescence was measured at 527 nm. The data was analyzed using the GraphPad prism 9.0 as described by Song *et al*. [32].

### Assay validation

The majority of HTS systems only allow for single testing of compounds, and therefore adequate sensitivity and high degree of accuracy are major requirements of the assays [30]. Assays for HTS also require reproducibility to accurately identify hit compounds among a very large number of compounds. After optimizing the different constituting parameters of the assay, the validation experiment was done by adapting the 3-day plate uniformity test. This uniformity test requires that for newly developed assays, the optimized assay conditions are tested for 3 consecutive days with no change in the conditions [42]. A total of five 384-well plates were used for the experiment by testing one plate each on day one and day 2 while three plates were tested on day 3 to understand the day-to-day and plate-to-plate variability of the assay. The interleaved-signal plate layout format was used for dispensing both positive and negative control into the wells. This means that the positive and negative controls were dispensed in alternate columns. Z’-factor was the statistical parameter used in the determination of the variability of the assay.

Z’-factor is a dimensionless and simple parameter which considers the difference in signal from samples and control as well as difference in the standard deviation of the controls. Z’-factor is used to assess the quality of the assay itself which makes use of the controls alone, therefore it was used in the validation of this assay [30]. 10 µL sample volume, mixture of CFP-PSB II (100 nM) and YFP-PAG (2 µM) was the negative control while the addition of 1 M GdnHCl to the mixture serves as the positive control.

A DMSO tolerance test was carried out on the assay to ascertain the amount of DMSO that the assay can withstand with no compromise to the FRET signal. Eight different DMSO concentrations were tested (0, 1, 1.5, 2, 3, 5, 7.5, and 10%).

### Compound screening

Peptidomimetics library of 15614 peptide-mimicking small molecule compounds (ChemDiv) was purchased from FIMM High Throughput Biomedicine Unit, University of Helsinki. Compounds were screened at 20 µM concentration with less than 1% DMSO. Compounds were dispensed with the ECHO acoustic liquid handler (Beckman Coulter). After which the FRET pair was dispensed with multichannel pipette to test for possible inhibition of the protein-peptide interaction.

Furthermore, both positive and negative controls were measured alongside the compounds to ascertain the Z’-factor value during screening. The fluorescence intensities at 477 nm and 527 nm were recorded with an Infinite Pro M1000 microplate reader (Tecan) using 430 nm as excitation wavelength. The apparent inhibition was calculated per cent of the PSB binding to the model peptide, where the negative control of the reaction was taken as 100 % binding and the positive control as 0 %.

### IC_50_ Measurement

For IC_50_ measurement, an increasing concentration of the compounds was added to the FRET pair. A log dilution series from 100 µM to 3.25 nM was considered for the compound concentration. The result was analyzed using GraphPad Prism 9.0. The FRET signal was plotted against the log of the inhibitor concentrations, and a nonlinear regression fit was used for the IC_50_ calculation.

### Differential scanning fluorimetry (DSF)

DSF measurements were carried out using Prometheus Nt.48 NanoDSF (NanoTemper Technologies GmbH, Munich, Germany) instrument. The melting temperatures (T_m_) were determined for both PSB domains with and without the selected hit compound OUL-PSBi-001. For the DSF analysis, the purified PSBs (without CFP-tag) [25,26] were diluted with either 20 mM Bis-Tris pH 6.8, 100 mM Glycine (PSB-I) or 20 mM Tris pH 8, 50 mM NaCl, 50 mM Glycine (PSB-II) to 10 µM. The analysis in the presence of the hit compound included 100 µM OUL-PSBi-001. The final DMSO concentration in all reactions was 1.3%. After preparation, samples were incubated at room temperature for 30 min for equilibration. Then the samples were loaded into standard capillary tubes, measurement was carried out from 20 °C to 90°C, with increment of 1°C per min. Five datasets were collected for each sample. An online tool MoltenProt [43,44] was used to calculate the T_m_ from NanoDSF data. Microsoft Excel (Microsoft Corporation, USA) was used to calculate the average values, and to draw the final figures.

### Bioluminescence based C-P4H activity assay

The C-P4H catalytic activity was measured with the insect-cell expressed full-length C-P4H-I and -II [8,33] and using a bioluminescent succinate detection assay (Succinate-Glo™, Promega), which quantifies the amount of succinate formed in the assay mixture from 2OG by C-P4H. Reaction mixtures (5 µL total volume) contain 5 µg/mL C-P4H, 100 µM peptide [(PPG)_10_], and a freshly prepared stock solution of 250 µM FeCl_2_, 2.5 mM ascorbate, and 500 µM 2OG (final concentrations in the assay mixture: 50 µM FeCl_2_, 500 µM ascorbate, and 100 µM 2OG) in 50 mM Tris buffer (pH 7.8). Enzyme and peptide were pre-incubated for 15 minutes at room temperature, and reactions were initiated by addition of the cofactor premix, followed by incubation for 30 minutes The reactions were then stopped by adding Reagent I (containing Succinate Buffer, Succinate-Glo Solution, and acetoacetyl-CoA), followed by a 1-hour incubation at 25 °C, allowing for the succinate dependent formation of ATP. Subsequently, reagent II, containing luciferase, was then added, and the amount of luminescence was measured after a 10-minute incubation time using a Spark microplate reader (Tecan). All measurements were performed in three independent experiments carried out on different days; error bars indicate the standard deviation between experiments.

### Docking of OUL-PSBi-001

Protein–ligand docking was performed using GOLD (CCDC, Cambridge, UK) [45] to investigate the mode of binding of OUL-PSBi-001 to the PSB domain. Docking was carried out using chain A of the PSB-I structure (PDB ID 9HT8) and chain A of the PSB-II structure (PDB ID 6EVN). To define the binding site, the peptide present in the structure (PPGPAGPPG) was extracted and used as a reference; the binding site was defined using a 10 Å radius around the peptide-defined pocket. Prior to docking, hydrogen atoms were added to the protein, and all non-protein atoms, including the peptide, crystallographic ligands, and water molecules, were removed. The ligand was prepared using Marvin (Marvin 19.1, ChemAxon). Docking was run using GOLD default settings, and poses were scored using the ChemPLP scoring function. Docking poses were inspected and visualized in Hermes (version 2021.3.0, Build 333819; CCDC, Cambridge, UK).

## Acknowledgements

We acknowledge funding from Biocenter Oulu spearhead project (LL) and Jane and Aatos Erkko Foundation (RW). The use of the facilities and expertise of the Biocenter Oulu Structural Biology core facility (a member of Biocenter Finland, Instruct-ERIC Centre Finland and FINStruct), Proteomics and Protein Analysis core facility (a member of Biocenter Finland) and Biocenter Oulu sequencing center are gratefully acknowledged. The assistance of Dr Albert Galera-Prat during the recombinant protein expression is gratefully acknowledged, and Professor Johanna Myllyharju and Dr Antti Salo for stimulating discussions.

## Author contributions

L.L. M.K.K. conceptualization; B.A., C.V.-R., S.M., M.M.R. investigation; B.A., C.V.-R. writing-original draft; All authors, writing, reviewing, and editing; L.L. and R.W. funding acquisition; C.V.R., R.W., L.L., M.K.K. supervision.

## Declaration of interests

The authors declare no competing interests.

## Additional information

All data generated or analysed during this study are included in this published article. The raw data generated during the current study are available from the corresponding authors on reasonable request.

## Notes

### Competing Interest Statement

The authors have declared no competing interest.

